## Supplementary Information Appendix 1 for "Gentle and fast all-atom model refinement to cryo-EM densities via Bayes’ approach"

### Supporting Information S1 Appendix

(Dated: February 28, 2023)

#### I. DERIVATION OF RELATIVE ENTROPY POTENTIAL FROM POISSON NOISE ASSUMPTION

By using Boltzman inversion for the total likelihood,

$$\log p\left(\rho|\rho^s(\vec{x})\right) = \frac{k}{k_B T} S(\rho, \rho^s(\vec{x})) + c\,. \tag{1}$$

and the assumption that electrons scatter independently in voxels

$$p(\rho|\rho^s(\vec{x})) = \prod_{v \in \text{voxels}} p(\rho_v|\rho^s_v) \tag{2}$$

$$\log p(\rho|\rho^s(\vec{x})) = \sum_{v \in \text{voxels}} \log p(\rho_v|\rho^s_v)\,, \tag{3}$$

as well as a Poisson distribution in each voxel with distribution parameters $\mathrm{Pois}(N\rho^s_v, \rho_v)$, where $\rho_v$ represents an interaction count,

$$p(\rho_v|\rho^s_v) = (N\rho^s_v)^{\rho_v} \exp(-N\rho^s_v)/\rho_v! \tag{4}$$

$$\log p(\rho_v|\rho^s_v) = \rho_v \log\left(N\rho^s_v\right) - N\rho^s_v - \log(\rho_v!)\,. \tag{5}$$

we obtain the similarity measure

$$\frac{k}{k_B T} S(\rho, \rho^s) = \sum_{v \in \text{voxels}} -N\rho^s_v + \rho_v \log N + \rho_v \log \rho^s_v - \log \rho_v!\,. \tag{6}$$

With the Stirling approximation $\log \rho_v! \approx \rho_v \log \rho_v - \rho_v$, assuming $\rho_v$ is sufficiently large, this can be simplified to

$$\frac{k}{k_B T} S(\rho, \rho^s) = -\sum_{v \in \text{voxels}} N\rho^s_v + \sum_{v \in \text{voxels}} \rho_v(\log N + 1) + \sum_{v \in \text{voxels}} \rho_v \log \rho^s_v - \rho_v \log \rho_v \tag{7}$$

According to eq. (1), similarity scores may be rescaled up to a constant which means that the term $\sum_v \rho_v(\log N+1)$ that does not depend on the model coordinates may be dropped from the right hand side of eq. (7). When the summed electron scattering probability is constant, e.g., when generating the model density from a sum of Gaussians as used below, the sum $\sum_v N\rho^s_v$ is also constant. With this regauging we get

$$\frac{k}{k_B T} S(\rho, \rho^s) = \sum_{v \in \text{voxels}} \rho_v \log \frac{\rho^s_v}{\rho_v}\,. \tag{8}$$

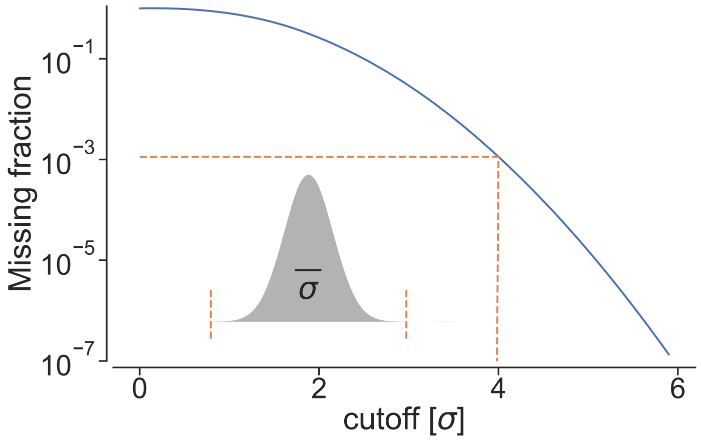


FIG. 1. Missing contribution from three-dimensional Gaussian mass density due to cutoffs at different $\sigma$. Orange lines indicate a cutoff at $4\sigma$, as used in all refinements in this publication.

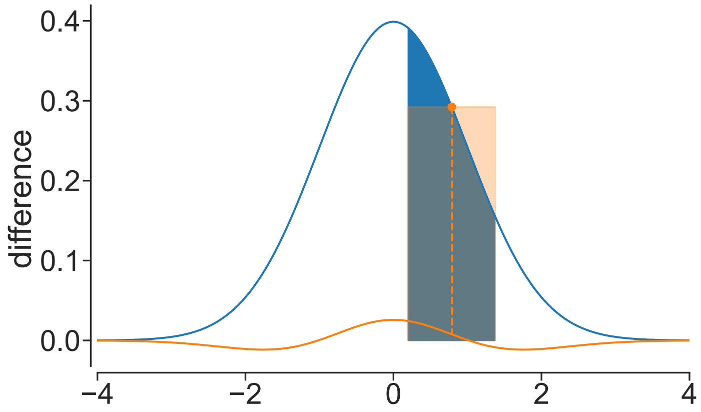


FIG. 2. Difference between integral over a voxel (blue area) of Gaussian function (blue line) and approximation by mid-point evaluation (orange) using a voxel size of $\sqrt{\log 4}\sigma$.

#### II. GAUSSIAN SPREADING ON A GRID

To determine the contribution of an atom at coordinate $r$ with an intensity $A$ to the model density, we project a Gaussian at $r$ onto the discrete grid used to describe cryo-EM densities by integration of the Gaussian over the voxel volume,

$$\rho_{\mathbf{v}} = \int_{x\in\mathbf{v}} A \frac{1}{\sqrt{2\pi}^3 \sigma^3} \exp\left[-\frac{(\mathbf{r}-\mathbf{x})^2}{2\sigma^2}\right] \tag{9}$$

Two approximations speed up computation of these contributions in practice. First, we ignore contributions to voxels with their centers further away than $4\sigma$ from the atom, since this overall amounts to very little loss of atom density as shown in Fig. 1. Second, we approximate the integral in eq. (9) by a discrete Gauss transform. Figure 2 shows the difference of the Gauss transform to integrating over voxel density when applying the $\sigma = \delta/\sqrt{\log 4}$ discretisation criterion as suggested in the main text.

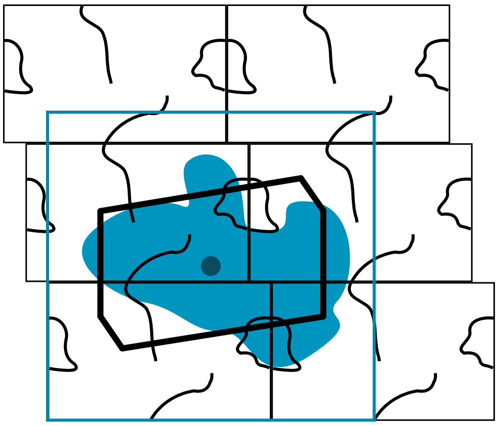

FIG. 3. When refining against a density (blue), only the periodic image of the atoms of the simulated molecules (black lines) is chosen that is closest to the center (blue dot) of the density boundary box (blue lines), effectively generating a Wigner-Seitz cell (thick black lines) of effective density influence around the density boundary box center.

#### III. THE MODEL DENSITY GRADIENT

Using the discrete Gauss transform on the voxel grid, the density spreading contributes to the force with the gradient of the model density,

$$\nabla_r \rho^{\mathrm{s}}_v(\mathbf{r}) = \sum_i -A_i \frac{(\mathbf{r_i}-\mathbf{v})}{\sigma} \frac{1}{\sqrt{2\pi}^3 \sigma^3} \exp[-\frac{(\mathbf{r_i}-\mathbf{v})^2}{2\sigma^2}] \,. \tag{10}$$

#### IV. EXPLICIT SIMILARITY MEASURE DEFINITIONS

The similarity measures over voxels $v$ are defined for inner-product, cross-correlation, swapped-relative-entropy and normal relative-entropy respectively as

$$S_{\mathrm{ip}}\left(\rho, \rho^{\mathrm{s}}\right) := \frac{1}{N_{\mathrm{voxel}}} \sum_v \rho_v \rho^{\mathrm{s}}_v \tag{11}$$

$$S_{\mathrm{cc}}\left(\rho, \rho^{\mathrm{s}}\right) := \frac{\sum_v \left[(\rho_v - \bar{\rho})(\rho^{\mathrm{s}}_v - \bar{\rho}^{\mathrm{s}})\right]}{\sqrt{\sum_v (\rho_v - \bar{\rho})^2 \sum_v (\rho^{\mathrm{s}}_v - \bar{\rho}^{\mathrm{s}})^2}} \tag{12}$$

$$S_{\mathrm{re-swapped}}\left(\rho, \rho^{\mathrm{s}}\right) := \sum_{v, \rho_v > 0, \rho^{\mathrm{s}}_v > 0} \rho^{\mathrm{s}}_v \log(\rho_v) \tag{13}$$

$$S_{\mathrm{re}}\left(\rho, \rho^{\mathrm{s}}\right) := \sum_{v, \rho_v > 0, \rho^{\mathrm{s}}_v > 0} \rho_v [\log(\rho^{\mathrm{s}}_v) - \log(\rho_v)] \tag{14}$$

#### V. TREATMENT OF PERIODIC BOUNDARY CONDITIONS

In the molecular dynamics simulation periodic boundary conditions are used. When multiple periodic images of the same atom are within the extent of the cryo-EM den-

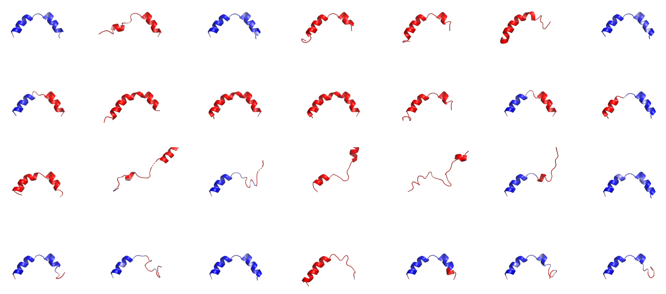

FIG. 4. Final results of helix to synthetic density refinement using adaptive force scaling from the aligned starting position using inner-product, cross-correlation, relative-entropy swapped, and relative entropy (ordered top to bottom).

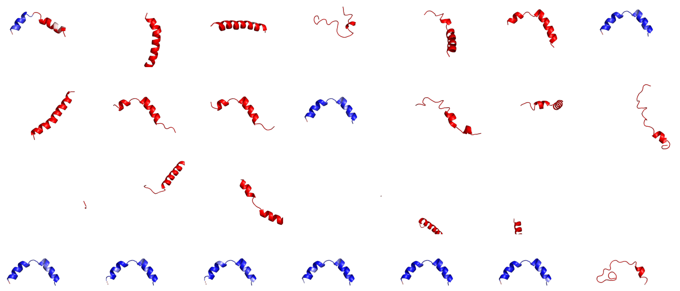

FIG. 5. Final results of helix to synthetic density refinement using adaptive force scaling from the unaligned starting position using inner-product, cross-correlation, relative-entropy swapped, and relative entropy (ordered top to bottom).

sity voxel grid (Fig. 3), only the one closest to the density map center is contributing to the model density. This allows refinement simulations without any restrictions on the respective size of simulation boxes and voxel grids and easy treatment with pressure coupling algorithms that change the size of the simulation box. In a few special setups, a situation may occur where different periodic images of the same molecule contribute to density based potential, thus "pulling" the molecule apart, which is easily avoided by either increasing the simulation box size or reducing the voxel grid, respectively.

#### VI. HELIX REFINEMENT

Figures 4 and 5 show the helix refinement result of all replicates starting from the aligned and unaligned positions for inner-product, cross-correlation, relative-entropy swapped, and relative-entropy based potentials.

TABLE I. Heavy-atom RMSD [Å] of helix fitting from the aligned starting position using inner-product, cross-correlation, relative-entropy swapped, and relative entropy at the final frame, as compared to the ground truth model.

| replicate | 1 | 2 | 3 | 4 | 5 | 6 | 7 |
|---|---|---|---|---|---|---|---|
| inner-product | 1.01 | 5.42 | 0.753 | 7.33 | 7.55 | 7.89 | 0.53 |
| cross-correlation | 3.96 | 6.41 | 5.86 | 5.92 | 7.32 | 3.85 | 3.93 |
| relative-entropy-swapped | 5.5 | 15.7 | 7.31 | 17.0 | 13.1 | 11.3 | 0.747 |
| relative-entropy | 2.07 | 4.21 | 0.732 | 7.19 | 2.06 | 2.38 | 3.55 |

TABLE II. Heavy-atom RMSD [Å] of helix fitting from the unaligned starting position using inner-product, cross-correlation, relative-entropy swapped, and relative entropy at the final frame, as compared to the ground truth model.

| replicate | 1 | 2 | 3 | 4 | 5 | 6 | 7 |
|---|---|---|---|---|---|---|---|
| inner-product | 4.05 | 23.3 | 12.2 | 12.5 | 22.4 | 7.83 | 0.832 |
| cross-correlation | 16.7 | 8.78 | 8.81 | 0.777 | 25.6 | 21.4 | 27.7 |
| relative-entropy-swapped | 50.2 | 33.2 | 26.8 | 56.7 | 62.2 | 43.5 | 74.0 |
| relative-entropy | 0.837 | 0.748 | 0.803 | 0.661 | 0.685 | 0.726 | 8.21 |

#### VII. STEREOCHEMISTRY AND MODEL STATISTICS FOR REFINEMENT AGAINST EXPERIMENTAL DATA

To assess the structural quality of the refined aldolase and GroEl subunit models, model statistics were calculated with PHENIX (version 1.18.2-3874), using MOLPROBITY, CABLAM, as well as EMRINGER. The model statistics reflect the stereochemical properties of the final refinement simulation frame without any further geometry optimization.

#### VIII. ALDOLASE REFINEMENT - LENGTH-SCALE DEPENDENT REFINEMENT QUALITY

Figures 6,7 show the difference in Fourier shell correlation between the manually built model and different density-guided simulation results. Positive values point to a fit that is better than the manually built model for a given length scale.

TABLE III. Heavy-atom RMSD [Å] of aldolase fitting using inner-product, cross-correlation, relative-entropy swapped, and relative entropy at the final frame, as compared to PDB id 6V20.

| replicate | 1 | 2 | 3 |
|---|---|---|---|
| inner-product | 0.967 | 0.871 | 0.926 |
| cross-correlation | 0.926 | 0.916 | 0.960 |
| relative-entropy-swapped | 0.908 | 0.924 | 0.937 |
| relative-entropy | 0.974 | 0.987 | 1.050 |

TABLE IV. Model statistics of PDB ids 6V20 and 6ALD

| | deposited model | starting model |
|---|---|---|
| PDB id | 6V20 | 6ALD |
| Ramachandran outliers (%) | 0 | 0 |
| favored (%) | 98.24 | 94.35 |
| Rotamer outliers (%) | 0.39 | 16.34 |
| C-beta deviations | 0 | 2 |
| Clashscore | 4.83 | 15.61 |
| RMS (bonds) | 0.0097 | 0.0049 |
| RMS (angles) | 2.30 | 1.09 |
| MolProbity score | 1.25 | 3.00 |

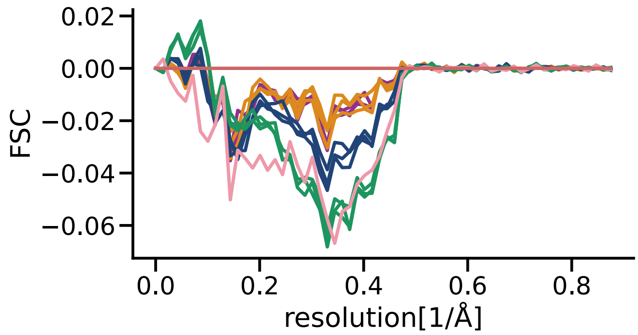


FIG. 6. Aldolase refinement. Difference in FSC to manually built model for inner-product (purple), cross-correlation (ochre), swapped relative-entropy (blue) and relative-entropy (green) based refinements final frame FSC (solid).

#### IX. GROEL REFINEMENT- LENGTH-SCALE DEPENDENT REFINEMENT QUALITY

Deviation from the published model for GroEl is shown in Table VI and Fig. 8. The reason for the worse performance of "relative-entropy" based potential is shown

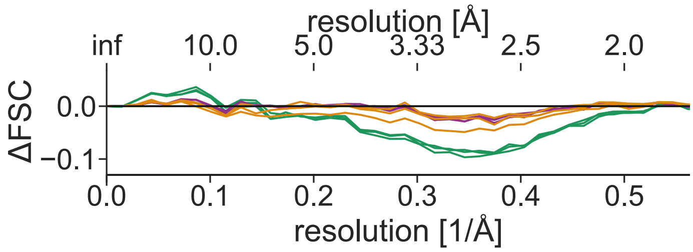


FIG. 7. Aldolase refinement from distorted structure. Difference in FSC to manually built model for inner-product (purple), cross-correlation (ochre), relative-entropy swapped (dark blue) and relative-entropy (green) based refinements with best accepted FSC (dotted) and final FSC (solid).

TABLE V. Model statistics of final frame of aldolase simulations.

| method | inner-product | | | cross-correlation | | | relative-entropy-swapped | | | relative-entropy | | |
|---|---|---|---|---|---|---|---|---|---|---|---|---|
| replicate | 1 | 2 | 3 | 1 | 2 | 3 | 1 | 2 | 3 | 1 | 2 | 3 |
| Ramachandran outliers (%) | 1.03 | 0.81 | 1.25 | 1.03 | 1.10 | 1.03 | 0.66 | 1.25 | 1.17 | 1.69 | 1.69 | 1.61 |
| Ramachandran favored | 96.85 | 96.26 | 96.48 | 96.19 | 95.31 | 96.19 | 95.82 | 95.23 | 95.82 | 95.01 | 94.65 | 93.70 |
| Rotamer outliers | 3.49 | 3.29 | 2.81 | 3.29 | 3.20 | 3.68 | 4.75 | 3.29 | 4.55 | 5.14 | 5.43 | 7.27 |
| C-beta deviations | 91 | 93 | 99 | 110 | 107 | 125 | 107 | 97 | 93 | 115 | 99 | 133 |
| Clashscore | 0.58 | 1.21 | 0.82 | 0.92 | 1.26 | 1.41 | 0.92 | 1.41 | 1.31 | 1.31 | 0.92 | 1.21 |
| RMS(bonds) | 0.0376 | 0.0384 | 0.0377 | 0.0391 | 0.0384 | 0.0433 | 0.0362 | 0.0357 | 0.0352 | 0.0363 | 0.0368 | 0.0375 |
| RMS(angles) | 3.51 | 3.50 | 3.52 | 3.61 | 3.61 | 3.76 | 3.50 | 3.51 | 3.48 | 3.55 | 3.50 | 3.58 |
| MolProbity score | 1.30 | 1.48 | 1.33 | 1.43 | 1.56 | 1.56 | 1.58 | 1.60 | 1.65 | 1.74 | 1.70 | 1.91 |
| CABLAM disfavored (<5%) | 5.3 | 5.6 | 5.5 | 5.4 | 5.8 | 5.5 | 5.5 | 7.0 | 5.2 | 7.0 | 6.2 | 7.2 |
| CABLAM outlier (<1.5%) | 1.8 | 1.4 | 2.0 | 1.5 | 1.7 | 2.1 | 1.5 | 2.6 | 2.1 | 2.6 | 2.2 | 2.0 |
| CABLAM CA geometry outlier (<0.5) | 0.44 | 0.37 | 0.52 | 0.59 | 0.52 | 0.37 | 0.66 | 0.59 | 0.59 | 0.44 | 0.81 | 0.74 |
| EMRinger Score | 6.03 | 6.06 | 6.49 | 6.13 | 6.04 | 6.41 | 5.96 | 5.75 | 5.42 | 5.03 | 5.64 | 5.40 |

TABLE VI. Model statistics of final frame of aldolase simulations.

| | ip | | | cc | | | logd | | | re | | |
|---|---|---|---|---|---|---|---|---|---|---|---|---|
| | 1 | 2 | 3 | 1 | 2 | 3 | 1 | 2 | 3 | 1 | 2 | 3 |
| Ramachandran outliers | 0.96 | 0.57 | 0.38 | 1.53 | 0.00 | 0.57 | 3.07 | 1.34 | 2.30 | 2.30 | 0.77 | 1.15 |
| favored | 94.83 | 94.83 | 95.40 | 93.49 | 95.59 | 94.83 | 89.08 | 91.38 | 92.53 | 89.08 | 92.34 | 93.68 |
| Rotamer outliers | 5.91 | 2.69 | 4.57 | 7.53 | 5.91 | 6.72 | 15.05 | 11.83 | 7.53 | 6.99 | 4.03 | 5.38 |
| C-beta deviations | 51 | 40 | 52 | 49 | 51 | 46 | 67 | 54 | 47 | 56 | 54 | 51 |
| Clashscore | 1.83 | 1.31 | 1.57 | 1.83 | 1.04 | 1.18 | 5.35 | 1.96 | 3.13 | 0.26 | 0.78 | 0.91 |
| RMS(bonds) | 0.0400 | 0.0377 | 0.0408 | 0.0398 | 0.0378 | 0.0365 | 0.0388 | 0.0375 | 0.0364 | 0.0373 | 0.0361 | 0.0347 |
| RMS(angles) | 3.56 | 3.56 | 3.63 | 3.62 | 3.53 | 3.57 | 3.83 | 3.70 | 3.61 | 3.70 | 3.64 | 3.63 |
| MolProbity score | 1.89 | 1.54 | 1.72 | 2.04 | 1.70 | 1.82 | 2.76 | 2.28 | 2.24 | 1.81 | 1.68 | 1.75 |
| CABLAM disfavored (<5%) | 7.3 | 6.5 | 7.1 | 7.5 | 8.3 | 7.3 | 14.4 | 12.7 | 11.5 | 14.8 | 9.4 | 7.7 |
| CABLAM outlier (<1.5%) | 1.7 | 1.3 | 1.9 | 2.3 | 2.7 | 2.1 | 7.1 | 4.4 | 4.4 | 5.6 | 3.1 | 2.3 |
| CABLAM CA geometry outlier (<0.5) | 0.96 | 0.77 | 0.77 | 0.58 | 0.77 | 1.15 | 2.12 | 1.73 | 1.73 | 1.92 | 0.77 | 1.54 |
| EMRinger Score | 2.08 | 2.14 | 2.05 | 1.73 | 1.51 | 1.13 | 2.19 | 2.05 | 1.88 | 1.35 | 1.69 | 1.95 |

TABLE VII. Unaligned heavy-atom RMSD [Å] of refinement using distorted model aldolase structure with inner-product, cross-correlation and relative entropy at best allowed and final frame, as compared to the manual model (pdbid:6V20).

| | Final |
|---|---|
| Inner-product | 1.07 |
| Cross-correlation | 1.02 |
| Relative-entropy | 1.13 |

TABLE VIII. Heavy-atom RMSD [Å] of GroEL fitting using inner-product, cross-correlation, relative-entropy swapped, and relative entropy at the final frame, as compared to PDB id 6V20.

| replicate | 1 | 2 | 3 |
|---|---|---|---|
| inner-product | 1.36 | 1.39 | 1.41 |
| cross-correlation | 1.38 | 1.4 | 1.41 |
| relative-entropy-swapped | 1.48 | 1.47 | 1.49 |
| relative-entropy | 1.96 | 1.81 | 1.75 |

in Fig. 9.

#### X. SCATTERING CROSS SECTION *A*

We approximate the scattering cross sections at 150keV for common biomolecules as listed in Table IX.

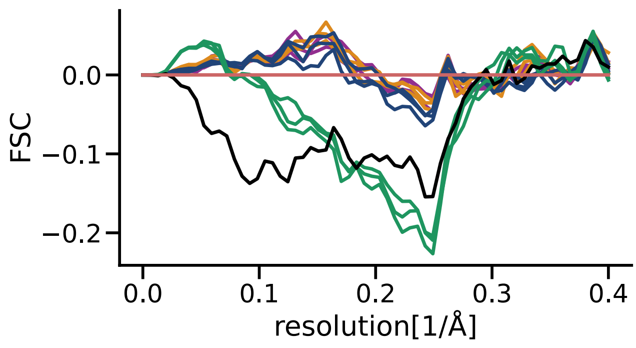


FIG. 8. GroEl subunit refinement. The difference in FSC to the deposited built model for inner-product (purple), cross-correlation (ochre), swapped relative-entropy (blue), and relative-entropy (green) based refinements. The simulation start is shown in black, published models in salmon.

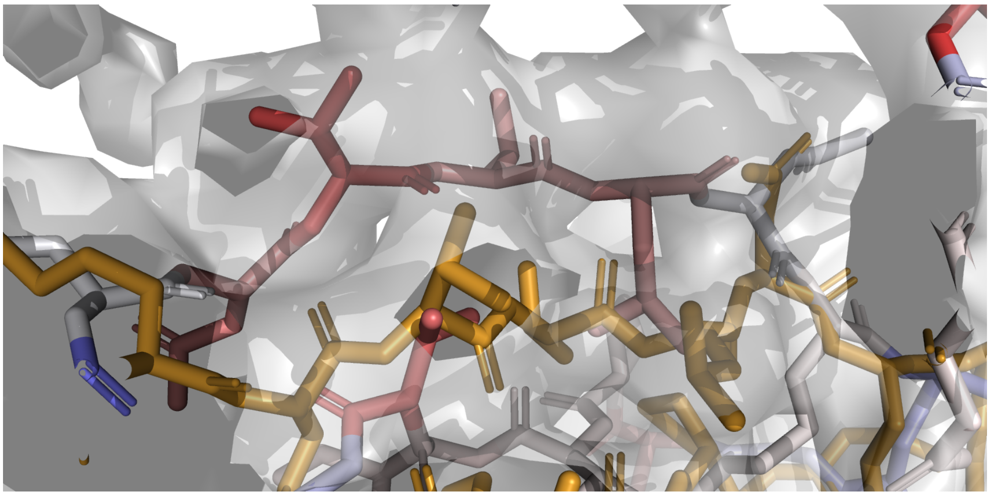

FIG. 9. Relative entropy based refinement (blue-red, according to RMSD to published model) struggles in regions with additional density, whereas the cross-correlation based potential (ochre) adheres to the local minimum defined by the density.

TABLE IX. Scattering cross section for common atoms at 150keV, from Appendix C in Ref. [1].

| Element | cross section | approximation |
|---|---|---|
| H | 0.04448 | 0 |
| C | 1 | 1 |
| N | 0.78061 | 1 |
| O | 0.62773 | 1 |
| P | 4.79207 | 1 |
| S | 4.24146 | 1 |

#### XI. ADAPTIVE FORCE SCALING - FORCE CONSTANT

Figure 10 shows the evolution of the adaptive force constant during helix refinement, starting from a heuristic value of 10 that guarantees minimal impact on the simulation from the start. The absolute value does not have any particular physical meaning or units since it's merely a relative factor describing the balance between force field and density fitting.

#### XII. REFERENCES

[1] Kirkland, E. J. *Advanced computing in electron microscopy* (Springer, 1998).

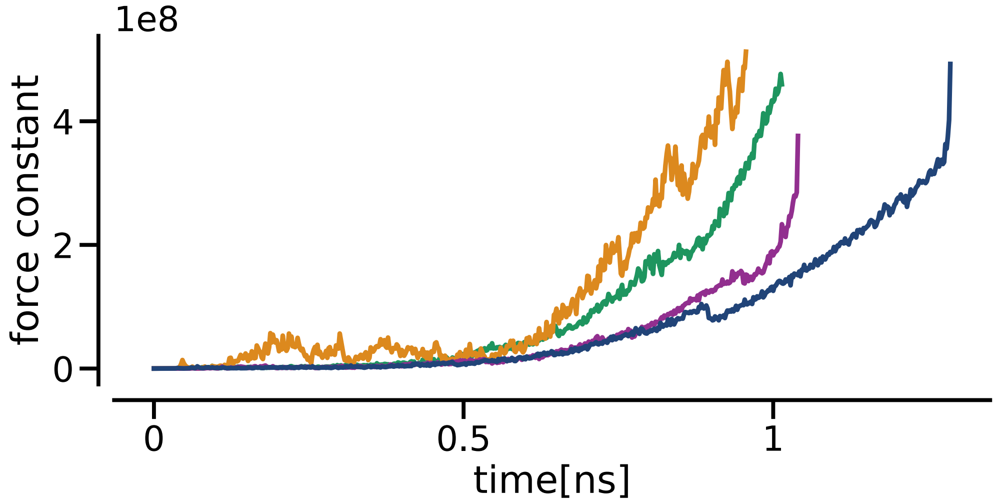


FIG. 10. Evolution of force constant during aligned helix refinement for inner-product(purple), cross-correlation(ochre), relative-entropy(green) and relative-entropy-swapped (blue).
